# The Evolution and Developmental Expression Profile of the *PIN-FORMED* Family in *Setaria viridis*

**DOI:** 10.1101/2025.10.07.680985

**Authors:** João Marcos Fernandes-Esteves, João Travassos-Lins, Juan David Ferreira Gomes, Marcio Alves-Ferreira

**Affiliations:** Universidade Federal do Rio de Janeiro (UFRJ), Laboratório de Genética Molecular e Biotecnologia Vegetal, Rio de Janeiro, Brazil

**Keywords:** Auxin transport, PIN-FORMED, Green foxtail, C4 grasses, Gene expression, Plant development

## Abstract

Auxin is one of the major driving forces of plant development and requires careful regulation of transporter proteins to establish polar auxin transport. The *PIN-FORMED* (*PIN*) family plays a pivotal role in plant development by establishing auxin gradients that govern vascular patterning and organogenesis. However, the *PIN* family remains severely underexplored in *Setaria viridis*, a well-established model for C_4_ monocots. In this study, we identified and characterized 13 *PIN* genes in the *S. viridis* genome. Phylogenetic and collinearity analyses revealed duplication events in the *SvPIN1, SvPIN5* and *SvPIN10* subfamilies. Structural analysis uncovered unique features, including potential pseudogenization of *SvPIN5a*. Expression profiling across five developmental stages unveiled the potential developmental roles of *SvPINs*, with *SvPIN1* and *SvPIN10* paralogues predominantly expressed in shoots and panicles, *SvPIN2* and *SvPIN9* in roots, while *SvPIN5b* showed leaf-enriched expression, suggesting potential involvement in Kranz anatomy development. Hormonal treatments in callus cultures revealed auxin-mediated upregulation of *SvPIN1b, SvPIN2, SvPIN5d, SvPIN8* and *SvPIN10a*. Our findings provide significant insights into the role of *PIN* genes in *S. viridis* and other C_4_ monocots, establishing a foundation for future functional studies and offering potential targets for crop improvement through auxin transport manipulation.

**Key Message:** This study characterizes the *PIN-FORMED* auxin transporter family in the C_4_ model *Setaria viridis*, revealing gene duplication, tissue-specific expression, and hormonal regulation that may influence C_4_ traits.

## Introduction

Auxin was the first ever discovered phytohormone (Darwin 1883) and was first isolated in the form of indole-3-acetic acid (IAA) (Went 1926). Throughout the decades of plant developmental science, it became clear that auxin is involved with a myriad of transcriptional and non-transcriptional signaling processes, influencing cell growth, organ initiation and patterning (Leyser 2018; Bogaert et al. 2022). One particularly important process regulated by auxin is vascular development. The evolution of vascular systems was pivotal for the colonization of terrestrial environments by land plants, enabling not only the rapid transport of molecules and signals, but also providing structural support (Ruonala et al. 2017). Vascular differentiation relies on the establishment of local auxin maxima and gradients, mediated by plasma membrane (PM)-localized transporter proteins in provascular cells (Petrášek and Friml 2009; Vosolsobě et al. 2020). The polar distribution of auxin transporters in the PM is essential for the unique directional flow of auxin, or polar auxin transport (PAT), which drives vascular bundle formation and leaf initiation (Scarpella et al. 2010). Beyond vascular development, PAT regulates embryogenesis, organogenesis, apical dominance and organ tropism (Ung et al. 2023).

Among the most well-studied auxin transporters is the *PIN-FORMED* (*PIN*) family. Since their discovery (Okada et al. 1991), PIN proteins have been recognized as central players in plant development. As efflux carriers, canonical PINs (PIN1, -2, -3/4/7, -10 and -11) are localized in the plasma membrane and export auxin to neighboring cells, establishing PAT and facilitating environmental and developmental responses (Adamowski and Friml 2015; Ung et al. 2023). Some PINs localize to the endoplasmic reticulum (ER) membrane, and thus are classified as non-canonical (PIN5, -6, -8 and -9), where they regulate intracellular auxin homeostasis by sequestrating auxin within the ER (Mravec et al. 2009; Ung et al. 2023).

PIN proteins are present in streptophyte algae, though their functions remain poorly understood and appear to differ significantly from those in land plants. Land plant *PINs* are thought to have originated from a single algal ancestor. (Viaene et al. 2013; Bennett 2015). *PIN* diversification coincides with that of land plants and key evolutionary innovations throughout their history. In bryophytes (non-vascular plants), PINs regulate critical developmental processes, including sporophyte development and spermatogenesis (Viaene et al. 2014; Lüth et al. 2023). During the rise of flowering plants, the *PIN-FORMED* family underwent major duplications, which contributed to floral development and the diversification of angiosperms (Zhang et al. 2020). Furthermore, *PIN* diversification has been linked to developmental innovations in monocots, particularly in grass architecture (Balzan et al. 2014; Wakeman and Bennett 2023), contributing to the major success of this group, which encompasses some of the world’s most important crops.

In addition to their role in plant development, especially in flowering and embryogenesis, *PIN* genes have been associated with abiotic stress responses. In sorghum, for instance, certain *PIN* family members colocalize with stay-green quantitative trait loci (QTLs) and have been associated with drought tolerance and increased yield (Borrell et al. 2022; Wong et al. 2023). Recent studies also implicate PIN proteins in the thermonastic movement (nondirectional responses to changes in temperature) of leaves mediated by PIN3 (Park et al. 2019), and changes in PIN2 polarization in response to oxidative stress (Zwiewka et al. 2019). Despite these advances, much remains unknown about auxin biology and the potential applications of *PIN* genes in crop improvement.

*Setaria viridis* is closely related to economically significant crops such as maize, sorghum, and sugarcane, sharing the same NADP-ME C_4_ photosynthetic pathway (Brutnell et al. 2010). This makes *S. viridis* an ideal model for genetic studies focused on C_4_ species. While *PIN-FORMED* genes have been extensively characterized in *Arabidopsis* and other C_3_ plants, their roles in C_4_ species remain largely unexplored. In this study, we systematically identified and characterized 13 *PIN-FORMED* genes in the *Setaria viridis* genome. Our integrative approach combined phylogenetic reconstruction, gene structure and promoter analyses, and developmental and hormonal expression profiling. We uncovered lineage-specific features such as potential pseudogenization of *SvPIN5a*, as well as tissue- and stage-specific roles for *SvPIN1, SvPIN2, SvPIN9*, and *SvPIN10* subfamilies.

## Materials and Methods

### Plant Materials

Seeds of the A10.1 accession (PI 669942 accession in the USDA Germplasm Resources Information Network) were germinated on filter paper following the gibberellic acid protocol described by Sebastian et al. (2014). Eight days after imbibition (DAI), seedlings were transplanted into a 3:1 (v/v) soil-vermiculite substrate and grown until flowering at 42 DAI. Plants were harvested for gene expression analysis at five key developmental stages, following the *S. viridis* BBCH scale proposed by Junqueira et al. (2020). The stages are marked by the full expansion of the first (1.10), third (1.30) and seventh (1.70) leaves, partial panicle emergence (5.50), and appearance of the first visible diaspores (7.10) (**Supplementary Fig. S1**).

For callus induction, dehusked seeds from the A10.1 accession were placed on callus induction medium (CIM) as described by Martins et al. (2019). Morphogenic calli were cultured for 15 days on media containing varying concentrations of the phytoregulators 2,4-dichlorophenoxyacetic acid (2,4-D) and Kinetin (K). Three experimental groups, each consisting of 15 calli, were grown on: standard CIM (2 mg/L 2,4-D + 0.5 mg/L K), Auxin-rich medium (3 mg/L 2,4-D + 0.25 mg/L K) and Cytokinin-rich medium (1 mg/L 2,4-D + 0.75 mg/L K). Three pools of five calli per treatment were harvested for molecular analysis.

### Orthologue Identification and Phylogenetic Analyses

Genomic data from 16 species (*Amborella trichopoda* v1.0, *Ananas comosus* v3, *Arabidopsis thaliana* TAIR10, *Brachypodium distachyon* v3.2, *Glycine max* Wm82.a4.v1, *Oryza sativa* v7.0, *Panicum virgatum* v5.1, *Physcomitrium patens* v3.3, *Populus trichocarpa* v4.1, *Selaginella moellendorfii* v1.0, *Setaria italica* v2.2, *Setaria viridis* v4.1, *Solanum lycopersicum* ITAG4.0, *Sorghum bicolor* v3.1.1, *Vitis vinifera* v2.1 and *Zea mays* RefGen_V4) representing major angiosperm lineages, lycophytes and mosses were retrieved from the Phytozome v13 database (Goodstein et al. 2012). *PIN-FORMED* orthologues were identified using HMMER v3.3.2 (http://hmmer.org; Finn et al., 2011) with the PIN protein HMM profile (PF03547) as a query. Adjusted HMM profiles were generated for each taxon, and results were manually curated via phylogenetic analysis, BLASTP (Altschul et al. 1990), and conserved domain analysis using InterProScan (Jones et al. 2014).

Amino acid sequences were aligned using MUSCLE in MEGA X (Kumar et al. 2018). A maximum-likelihood phylogenetic tree was constructed using IQ-TREE v2.0.7 (Minh et al. 2020), with branch support assessed using 2000 bootstrap replicates (Hoang et al. 2018) and 2000 single-branch test iterations (Guindon et al. 2010). The JTT+R7 amino acid substitution model was selected according to the Bayesian information criterion by the ModelFinder algorithm (Kalyaanamoorthy et al. 2017). The *Klebsormidium flaccidum PIN* orthologue (GenBank AJA79000.2), identified via BLASTP, served as an outgroup.

### Chromosomal Localization and Duplication Analysis

Chromosomal positions of *S. viridis PIN* genes were extracted from the Phytozome v13 annotation file and visualized using the chromoMap R package (Anand 2018). Gene duplication events were identified with MCScanX (Wang et al. 2012), and collinearity maps were generated using the AccuSyn online platform (Núñez Siri et al. 2020).

### Gene Structure, Conserved Motifs and Transmembrane Domain Analyses

Gene structure (intron/exon composition) were derived from the Phytozome v13 annotation and visualized with GSDS 2.0 (Hu et al. 2015). Conserved motifs in the amino acid sequence of *S. viridis* PIN proteins were identified using MEME v5.5.7 (Bailey and Elkan 1994). Transmembrane domain structure was predicted using DeepTMHMM v1.0.44 (Hallgren et al. 2022) and visualized using MembraneFold v0.0.71 (Gutierrez et al. 2022).

### Quantitative Gene Expression Analysis

Total RNA was extracted from frozen plant and callus tissue using the ReliaPrep RNA Tissue Miniprep System (Promega™), followed by cDNA synthesis with TaqMan Reverse Transcription Reagents (Thermo Scientific™). Primer design was performed using Primer3Plus (Untergasser et al. 2007). Reference *genes folyl-polyglutamate synthase* (*Si035045*) and *phosphoglucomutase* (*Si034613*) were used for normalization during expression analysis (Lambret-Frotté et al. 2015). Quantitative PCR was conducted on a 7500 Fast Real-Time PCR System (Applied Biosystems™) according to the Platinum Taq DNA Polymerase (Invitrogen™) protocol. Raw fluorescence data was processed with Real-Time PCR Miner (Zhao and Fernald 2005). Relative expression across developmental stages was analyzed using qBase+ v3.3 (Hellemans et al. 2008), while differential expression in callus was assessed with REST-384 (Pfaffl 2002).

### Cis-Acting Regulatory Elements (CARE) Analysis

The 2000 base pair upstream region of each *S. viridis PIN* locus was analyzed for CAREs using the New PLACE database (Higo et al. 1999). Identified CAREs were classified into discrete categories based on the database annotation. CARE enrichment analysis was performed in RStudio using a Fisher exact test with Benjamini-Hochberg false discovery rate (FDR) correction. Results were plotted using the *ggplot2* (Wickham 2016) R package.

## Results

### Phylogenetic Analysis Reveals 13 PIN-FORMED Genes in Setaria viridis

PIN-FORMED proteins underly plant evolution since the first land plants began colonizing the aerial environment. Understanding the evolutionary history of *PINs* is a crucial step in the characterization of the family. Therefore, extensive phylogenetic analysis identified 13 *PIN-FORMED* genes in the *S. viridis* genome (**Table 1**), distributed across seven subfamilies (**Fig. 1A; Supplementary Fig. S2**). The *PIN1* clade comprises four paralogues (*SvPIN1a, SvPIN1b, SvPIN1c* and *SvPIN1d*), divided into two subfamilies: *PIN1* (*SvPIN1a* and *SvPIN1b*) and *SISTER OF PIN1* (*SoPIN1*; *SvPIN1c* and *SvPIN1d*). These genes are scattered across chromosomes 4, 1, 7 and 8, respectively (**Fig. 1B**). Collinearity analysis revealed that segmental duplications gave rise to the *SvPIN1a/1b* and *SvPIN1c/1d* pairs (**Fig. 1C**). Only one *PIN2* subfamily member (*SvPIN2*) was identified, located on chromosome 4 (**Fig. 1B**), and was classified as a dispersed duplication event by the collinearity algorithm. Notably, *S. viridis* lacks genes from the *PIN3/4/7* clade, which is predominantly found in eudicots (**Fig. 1A**). Conversely, the *PIN10* subfamily, a monocot-specific sister clade to *PIN3/4/7* (**Fig. 1A**), consists of two paralogues (*SvPIN10a* and *SvPIN10b*), located on chromosomes 3 and 5, respectively (**Fig. 1B**), and likely arose from a segmental duplication (**Fig. 1C**).

**Table 1:**
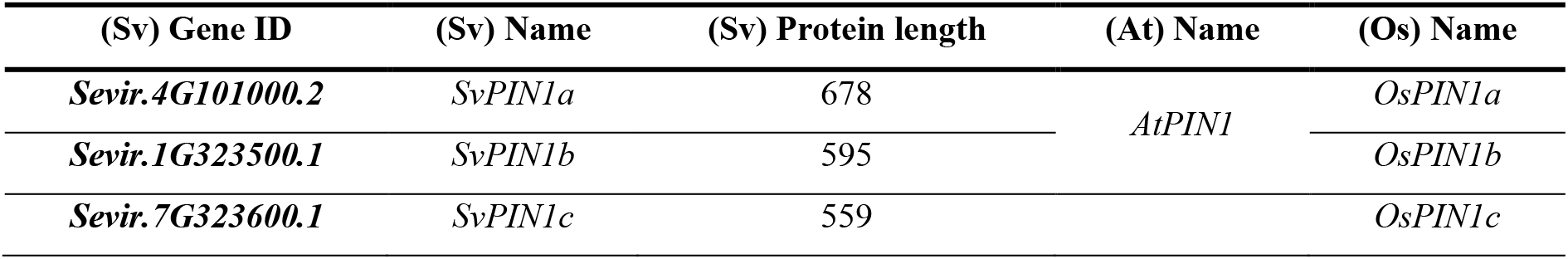
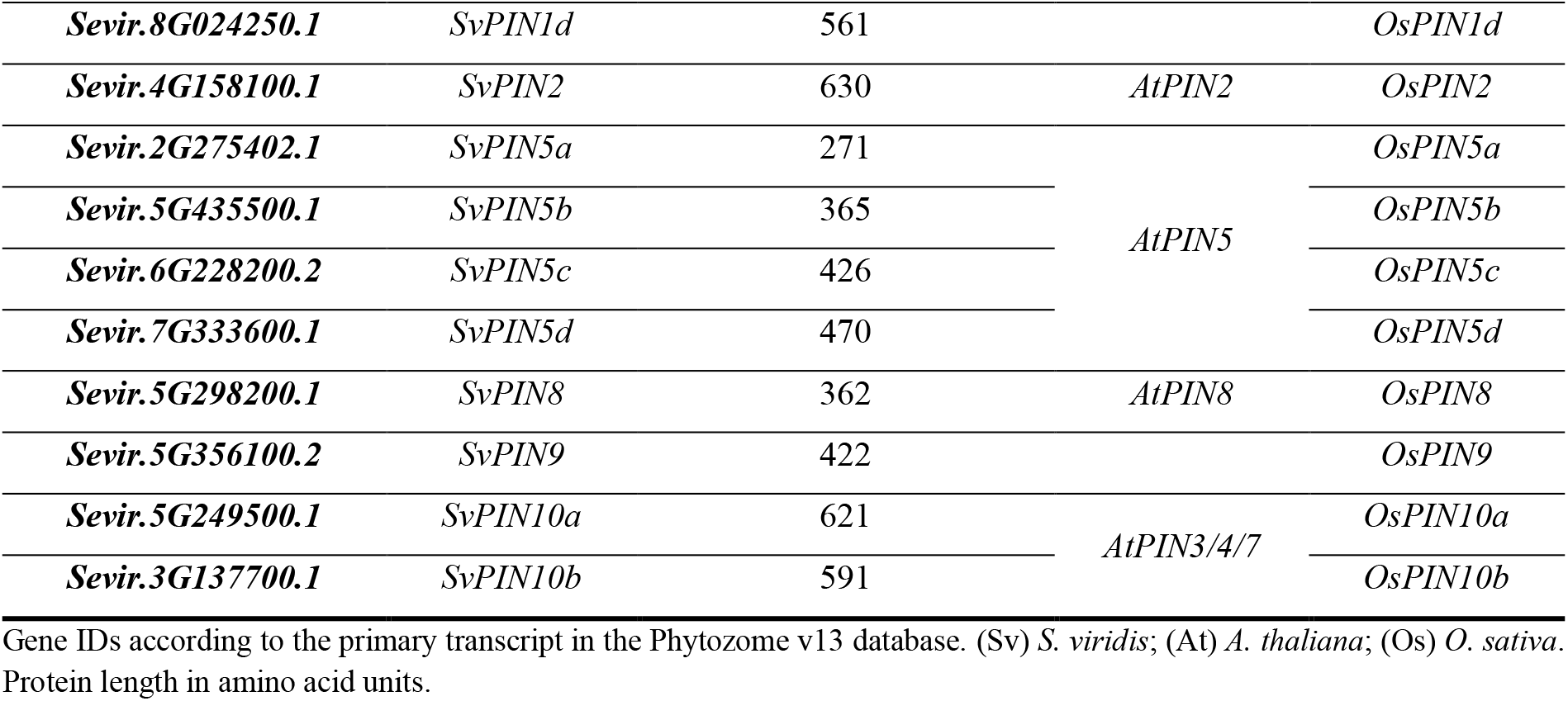
Identified PIN proteins of *Setaria viridis*.

**Fig. 1:**
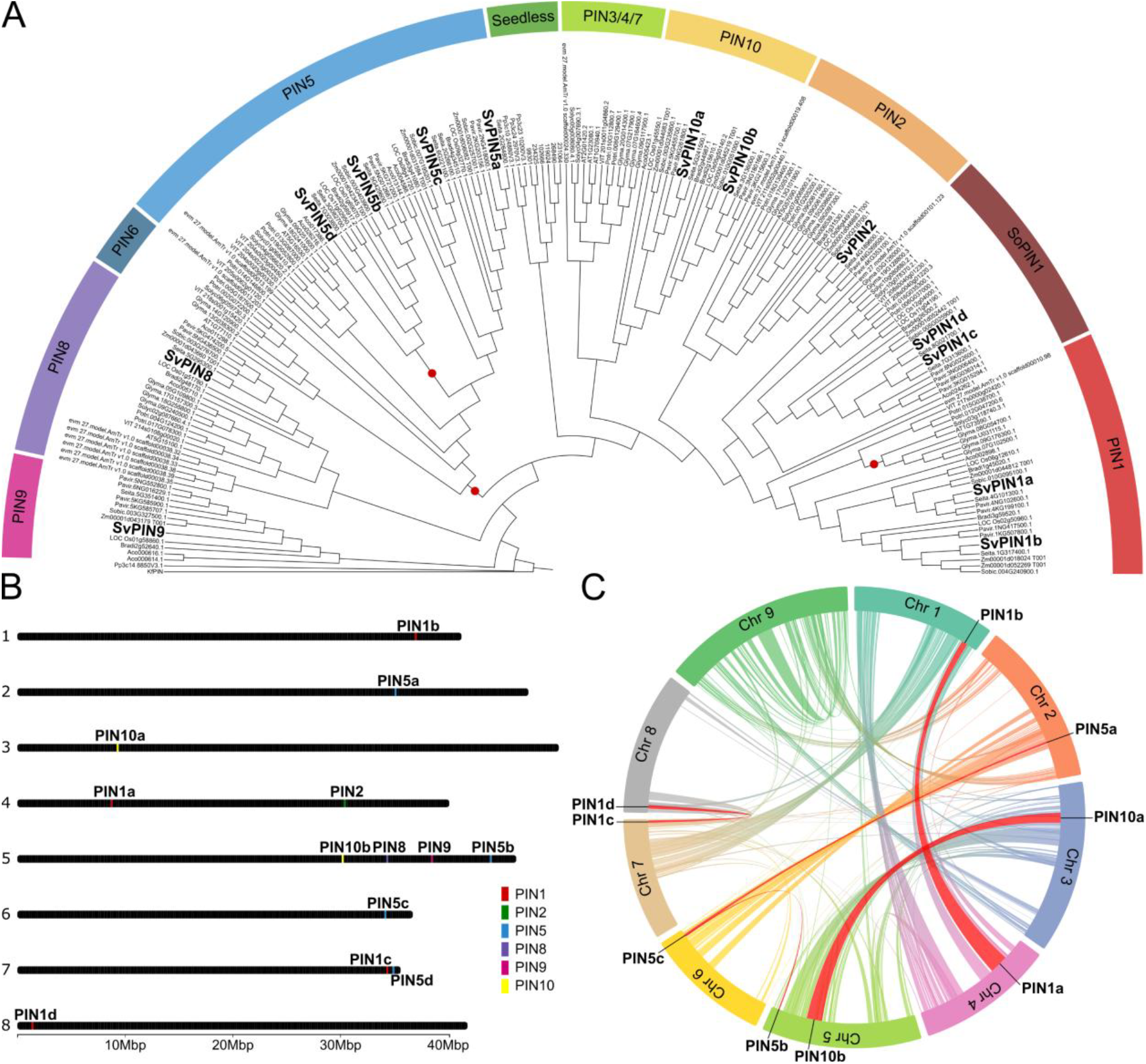
*Setaria viridis PIN-FORMED* family phylogenetic reconstruction, chromosomal location and syntenic analysis. **(A)** Maximum likelihood tree showcasing the evolutionary relationship of *PIN* subfamilies across land plants. The phylogeny encompasses representatives of mosses (*P. patens*), lycophytes (*S. moellendorfii*) and major groups of angiosperm species, focusing on grasses. *S. viridis* orthologues are highlighted in bold. Branches with support lower than 70% are marked with a red dot. The charophyte *K. flaccidum PIN* orthologue (*KfPIN*) was used as the outgroup. **(B)** Location of *SvPIN* genes in *S. viridis* chromosomes. Only the chromosomes containing *SvPIN* homologues are represented. **(C)** Collinearity map of *S. viridis* chromosomes and the identified syntenic blocks. *SvPIN1a, SvPIN1b, SvPIN1c, SvPIN1d, SvPIN5a, SvPIN5b, SvPIN5c, SvPIN10a* and *SvPIN10b* are represented within their respective duplicated regions. Duplicated regions are highlighted in red.

The *PIN5* subfamily includes four paralogues (*SvPIN5a, SvPIN5b, SvPIN5c* and *SvPIN5d*), located on chromosomes 2, 5, 6 and 7, respectively (**Fig. 1B**). *SvPIN5a, SvPIN5b* and *SvPIN5c* likely originated from segmental duplications (**Fig. 1C**), while *SvPIN5d* is a dispersed genomic duplicate. No orthologues of *PIN6* were identified in any of the grass genomes, including *S. viridis*. The *PIN8* and *PIN9* subfamilies each contain a single gene (*SvPIN8* and *SvPIN9*, respectively), both located on chromosome 5 (**Fig. 1B**). Collinearity analysis indicated that both genes originated from dispersed duplication events. Notably, the *PIN9* subfamily is another monocot-exclusive clade. Alongside the phylogenetic reconstruction of the *PIN-FORMED* family, we investigated the exon-intron composition of each gene and the conserved motifs in the amino acid sequences to corroborate the differences between canonical and non-canonical proteins.

### Gene Structure and Conserved Motifs Distinguish Canonical and Non-Canonical PINs

PIN-FORMED proteins are known for their highly conserved transmembrane domains, which are crucial for auxin transportation. Gene structure analysis showed that most *SvPIN* genes contain six exons, with a large initial exon, followed by a mid-sized exon and several smaller downstream exons (**Fig. 2A**). Interestingly, *SvPIN5a* lacks several downstream exons and encodes a significantly shorter transcript. When analyzing conserved protein motifs, we found high conservation across *S. viridis* PINs, particularly at their N- and C-termini, where the transmembrane helixes are located, while the center portion of the protein is more variable (**Fig. 2B**). Notably, the canonical PINs (PIN1, -2 and -10) present more conserved central motifs, while the non-canonical PINs (PIN5, - 8 and -9) exhibit less conservation. SvPIN5a was the only protein lacking the two C-terminal motifs that are present in all other family members, potentially due to its shorter transcript and likely compromising its tridimensional structure. To further corroborate the motif analysis, the transmembrane domains and protein structures were predicted using deep learning algorithms. These analyses revealed 10 highly conserved transmembrane helixes flanking the central hydrophilic loop in all proteins, except for PIN5a (**Fig. 2C-F**). Canonical PINs showed a larger hydrophilic loop than non-canonical PINs, reinforcing their identities. SvPIN9 has an intermediary-sized central loop (**Fig. 2E)**. Owing to the lack of the last two conserved motifs, SvPIN5a displayed an atypical predicted structure, with only seven transmembrane helixes (**Fig. 2F; Supplementary Fig. S3**). These combined results not only reinforce the divisions between canonical and non-canonical proteins but also point to SvPIN5a being functionally compromised. To investigate whether the 13 identified *PIN* genes are functional, gene expression analyses were performed throughout plant development and in callus cultures.

**Fig. 2:**
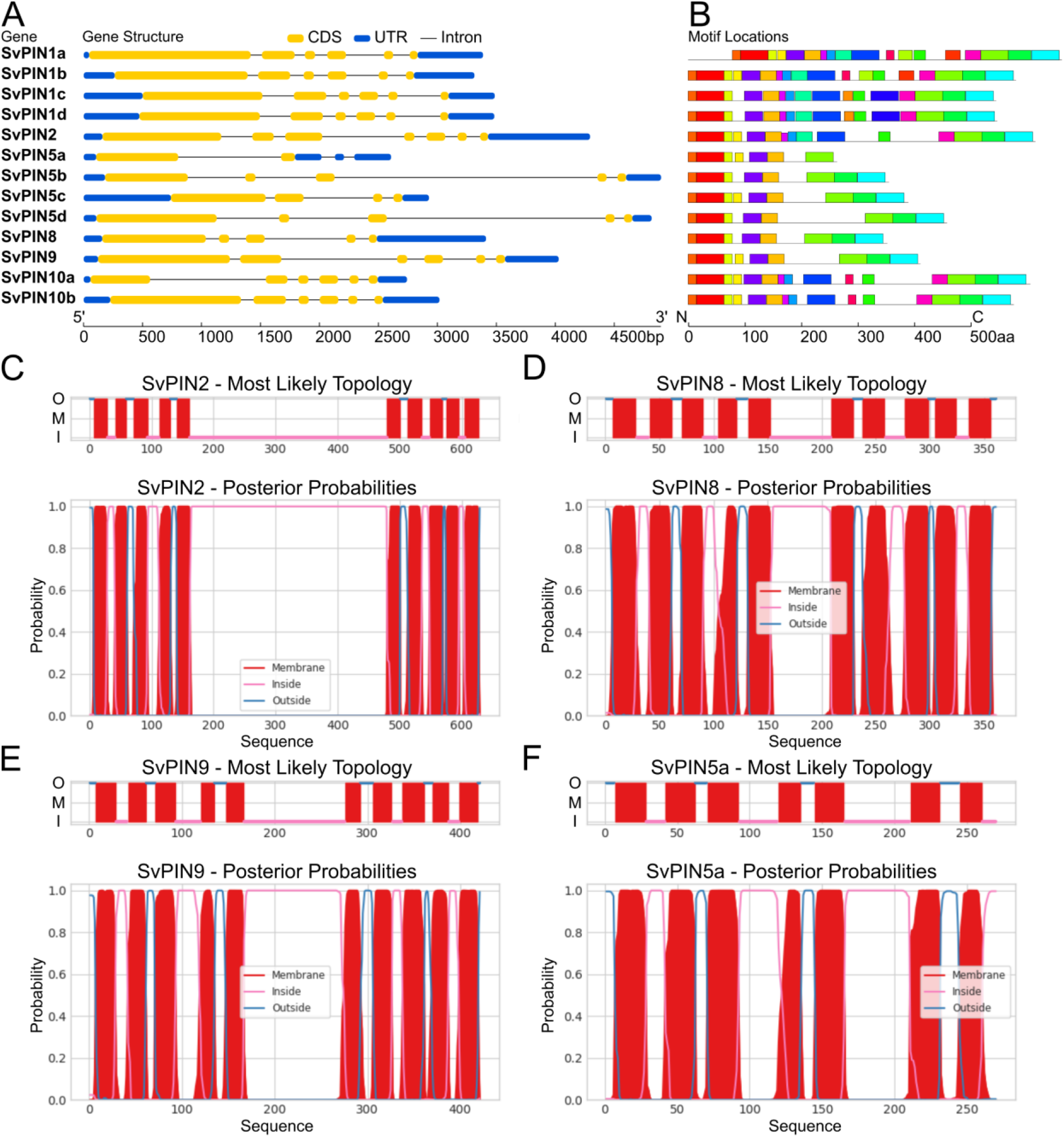
*Setaria viridis PIN-FORMED* gene structure, conserved motifs and transmembrane domain analyses. **(A)** *SvPIN* gene structure, showcasing the exon and intron distribution within the primary transcripts. Canonical PINs display transcripts with shorter introns, contrary to non-canonical PINs. *SvPIN5a* has a significantly smaller transcript and only two exons, reflected in its protein structure. **(B)** SvPIN proteins conserved motifs. Canonical PINs have more conserved motifs in the central hydrophilic loop portion of the protein compared to non-canonical PINs. The absence of the two C-terminal motifs in the SvPIN5a structure likely compromises its transporter function. The consensus sequence for each identified motif is available in **Supplementary Fig. S4. (C-F)** Transmembrane domain predictions for different PIN proteins. O = Outside; M = Membrane; I = Inside. **(C)** SvPIN2 and **(D)** SvPIN8 represent canonical and non-canonical proteins, respectively, displaying the significant size different between the hydrophilic loops of each group. **(E)** SvPIN9 showcases the intermediary-sized loop of a non-canonical protein derived from a canonical ancestor. **(F)** SvPIN5a lacks three transmembrane domains, possibly owing to the loss of the last two C-terminal conserved motifs that are present in every other PIN protein.

### Expression Profiling Highlights Tissue- and Stage-Specific Roles for SvPIN Genes

*PIN-FORMED* genes are shown to regulate a wide range of processes and therefore exhibit diverse expression patterns in multiple tissues and throughout all stages of development. To evaluate the role of PINs during *S. viridis* development, we analyzed five key developmental stages, from 8 days after imbibition (DAI) seedlings to 42 DAI flowering panicles (P1.10 to I7.10, Junqueira et al., 2020). The *SvPIN1* subfamily was mostly expressed in seedlings, shoots and panicles. Notably, all *SvPIN1* genes had very low expression in developed leaves (**Fig. 3A-D**). *SvPIN1a* was highly expressed in mature roots (R5.50) and shoots (C5.50), while *SvPIN1b* expression peaked during panicle development (I5.50). *SvPIN1c* was mostly expressed in shoot and panicle tissue throughout stages 1.70, 5.50 and 7.10. Similarly, *SvPIN1d* expression peaked in developing shoots (C1.70), and it was expressed twice as much as other *SvPIN1* genes in seedlings. *SvPIN2* was moderately expressed in seedlings and throughout all stages of root development (R1.30, R1.70 and R5.50) and peaked in flowering panicles (**Fig. 3E**). The *SvPIN5* subfamily (**Fig. 3F-H**) showed a peculiar expression profile. *SvPIN5a* expression remained undetected in all tissues and stages investigated, further pointing to a non-functional gene. Meanwhile, *SvPIN5b* was highly expressed in developed leaves (F1.30, F1.70 and F5.50), unlike any other *SvPIN*. Furthermore, *SvPIN5b* was moderately expressed in seedlings. *SvPIN5c* and *SvPIN5d* had similar expression profiles, being moderately expressed in seedlings and peaking during panicle development. Notably, *SvPIN5d* was the most expressed gene in flowering panicles. *SvPIN8* was mostly expressed in seedlings, shoots and panicles, while expression levels in roots and leaves remained low (**Figure 3I**). *SvPIN9* stood out for its peculiar expression profile (**Figure 3J**). While expression remained low in almost all tissues, *SvPIN9* had the highest expression peaks of all *SvPIN* genes in developing (R1.70) and mature roots (R5.50). Additionally, *SvPIN9* was moderately expressed in seedlings. Finally, the *SvPIN10* subfamily was mostly expressed in the reproductive tissues during panicle development and flowering (**Figure 3K-L**). *SvPIN10b* additionally showed high expression levels during shoot development. To test whether the observed *SvPIN* expression patterns are hormone-sensitive, we applied 2,4-D and kinetin treatments to callus cultures and compared the hormonal regulation of *SvPIN* genes with their developmental profiles.

**Fig. 3:**
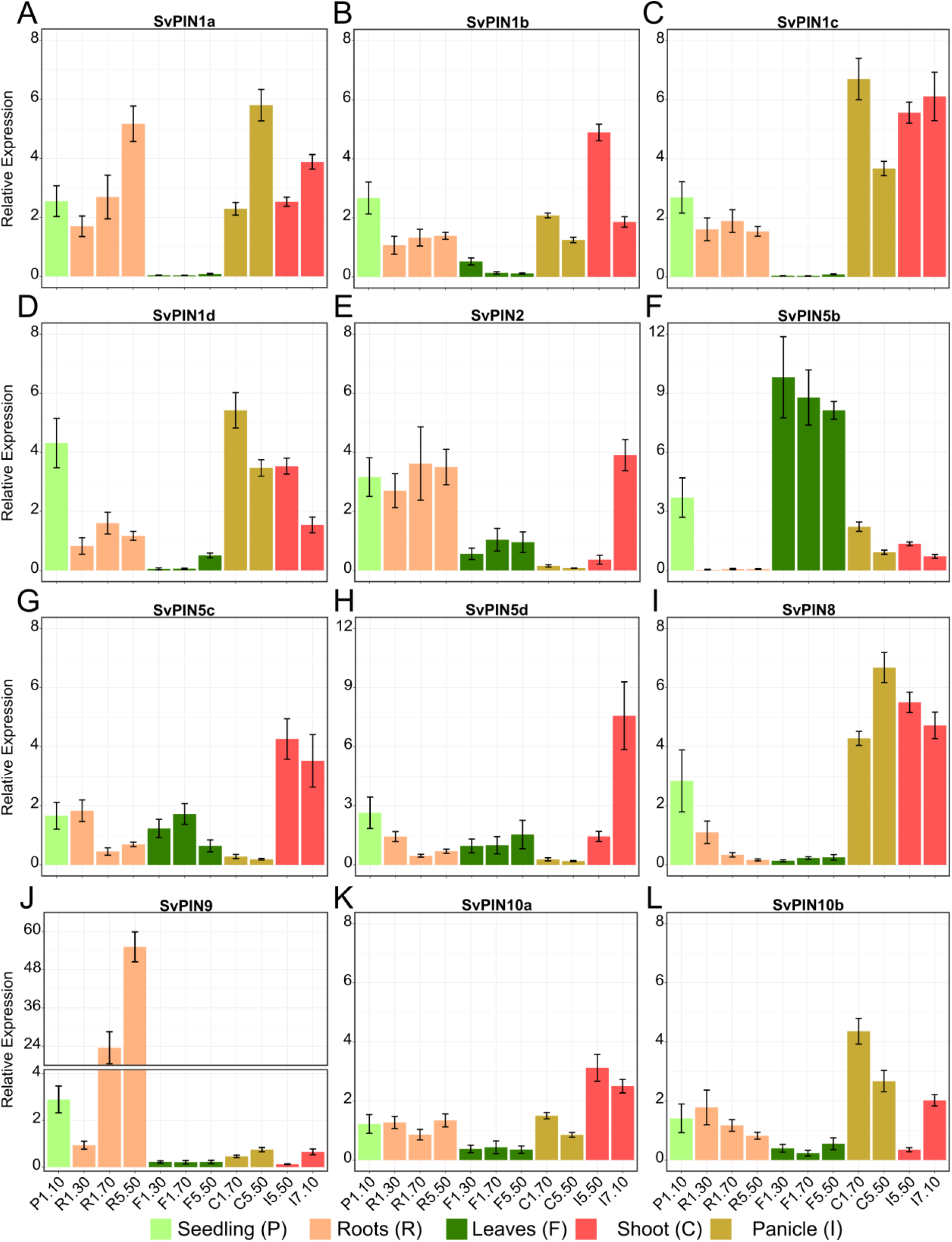
Relative expression of *Setaria viridis PIN-FORMED* genes throughout development. **(A-D)** *SvPIN1* subfamily gene expression profile. **(E)** *SvPIN2* gene expression profile. **(F-H)** *SvPIN5* subfamily gene expression profile. *SvPIN5a* expression was undetected in all tissues. **(I)** *SvPIN8* gene expression profile. **(J)** *SvPIN9* gene expression profile. Y-axis was cut to fit the relative expression bars of R1.70 and R5.50 samples. **(K-L)** *SvPIN10* subfamily gene expression profile. Expression values are represented as relative expression in comparison to reference genes *Si035045* and *Si034613*. Each bar represents an average between three biological replicates and error bars represent the standard deviation of each sample. P = Seedling; R = Roots; F = Leaves; C = Shoot; I = Panicle.

### Phytohormone Treatments Reveal Differential Regulation of SvPIN Genes in Calli

Plant calli, when induced by exogenous phytohormones, demonstrably develop into vascular-rich masses of undifferentiated tissue (Sugimoto et al. 2010), thus being interesting targets for the study of *PIN-FORMED* genes. *SvPIN* expression was responsive to different levels of 2,4-D and Kinetin during callus growth. The control calli rarely developed leaf-like structures and remained mostly unchanged during the course of the experiment (**Fig. 4A**). Calli treated with higher 2,4-D and lower Kinetin frequently developed pale, leaf-like structures, and often achieved larger mass than others (**Fig. 4B**). *SvPIN1b, SvPIN2, SvPIN5d, SvPIN8* and *SvPIN10a* were significantly upregulated in 2,4-D-treated calli (**Fig. 4D**). None of the *PIN-FORMED* genes were downregulated under high 2,4-D treatment. Conversely, low 2,4-D and high Kinetin levels resulted in calli developing fine, hair-like structures on the surface, reminiscent of young roots (**Fig. 4C**). Nearly all *SvPIN* genes had their expression repressed by this treatment, with the exception of *SvPIN8. SvPIN2, SvPIN5c* and *SvPIN5d* were the most downregulated genes (**Fig. 4D**). *SvPIN5a* expression remained undetected in the callus assays, corroborating the pseudogenization hypothesis. This analysis shows the hormone-sensitivity of *SvPINs* and how high concentrations of auxin lead to increased *PIN* expression and leaf initiation in undifferentiated tissue. To further support the expression data, we analyzed the cis-element composition of *PIN* promoters in *S. viridis*.

**Fig. 4:**
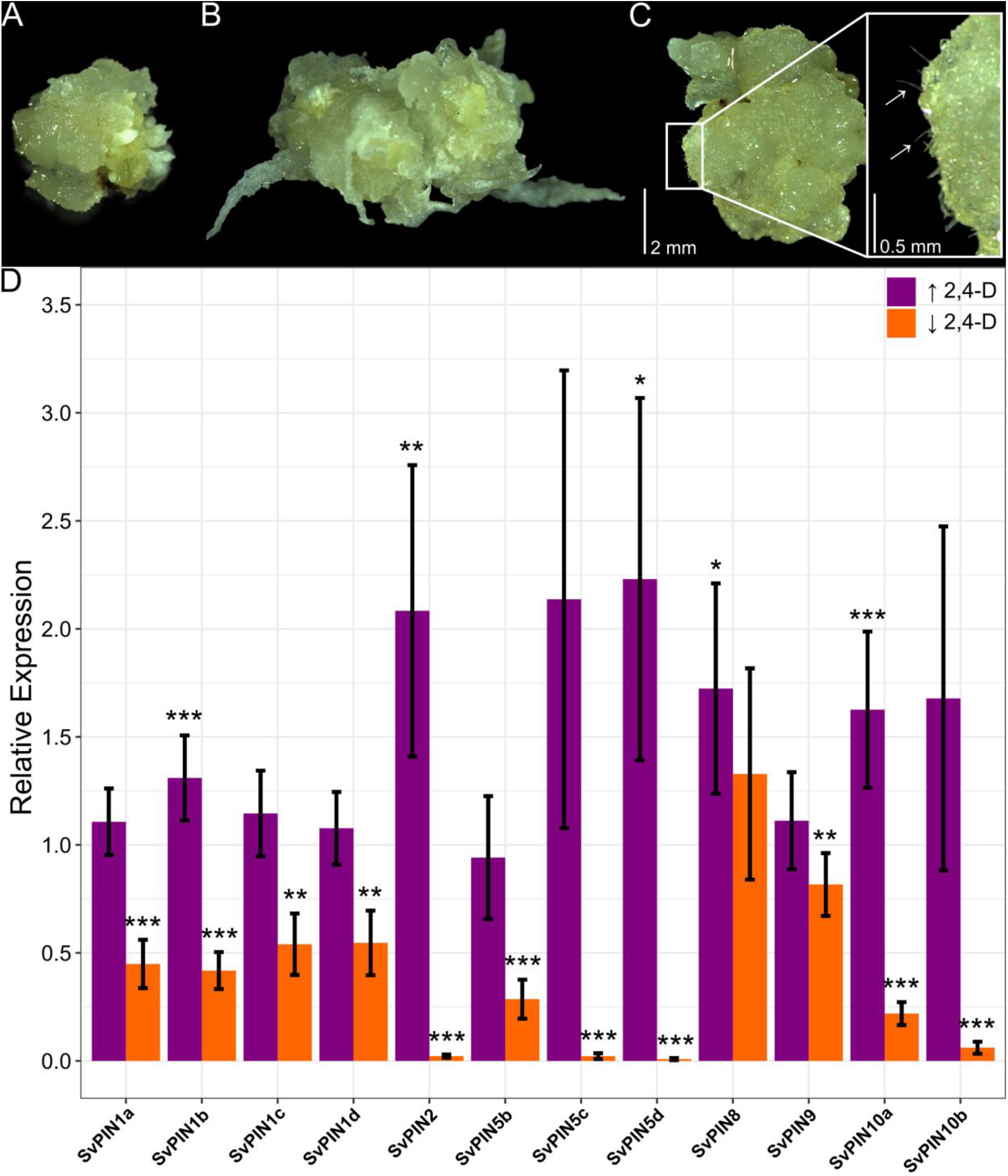
Differential expression of *Setaria viridis PIN-FORMED* genes in morphogenic callus under different 2,4-D and Kinetin concentrations. **(A)** Control morphogenic callus, cultivated in standard CIM. **(B)** Morphogenic callus under high 2,4-D, low Kinetin. Compared to control calli, these grew larger and developed leaf-like structures on their surface. **(C)** Morphogenic callus under low 2,4-D, high Kinetin. These retained a similar size compared to control calli, and developed fine, hair-like structures, similar to young roots, indicated by the arrows. **(D)** Relative expression of *SvPIN* genes under high (↑2,4-D) and low (↓2,4-D) concentrations of 2,4-D, respectively. Expression values are represented as relative expression in comparison to control expression levels and reference genes *Si035045* and *Si034613*. Each bar represents an average between three biological replicates and error bars represent the standard deviation of each sample. Asterisks represent significance according to the fixed reallocation randomization test, with P < 0.05 (*), P < 0.01 (**), P < 0.001 (***).

### Promoter Analysis Reveals a Wide Range of CAREs in SvPIN Genes

Cis acting regulatory elements (CAREs) are important aspects of gene expression and can offer insights into the potential roles of gene when paired with expression data. Promoter analysis identified 200 unique CAREs across all *SvPIN* genes (**Table S1**), categorized into eight functional groups: (1) Stress Response; (2) Light Response; (3) Tissue Expression; (4) Phytohormone Response; (5) Nutrient Response; (6) Defense Related; (7) Anaerobic Response; and (8) Others. CARE distribution was largely homogenous, though duplicate pairs, such as *SvPIN1a/1b* and *SvPIN1c/1d*, showed slightly divergent promoter compositions. In contrast, *SvPIN5a/5b* promoters shared remarkable similarities in CARE composition (**Fig. 5**). CARE enrichment analysis identified 20 significantly enriched CAREs in some promoters when compared to the rest of the family (**Table S2**). The *SvPIN1d* promoter was enriched in light-responsive motif S000486, linked to photomorphogenesis. *SvPIN5b* promoter was enriched in S000277, a motif related to endosperm-specific expression. *SvPIN5d* had the highest number of enriched motifs, including light response and defense related S000042, tissue expression-related elements S000512 and S000314 associated with root hairs and leaf expression, respectively, nutrient-responsive elements S000501 and S000507, and dehydration response elements S000414, S000415 and S000407. The *SvPIN9* promoter was enriched in phytohormone-responsive element S000390, linked to salicylic acid response. Finally, *SvPIN10a* was enriched in S000254, a tissue expression-related element associated with expression in pollen grains. These results align with some of the observations derived from expression data, further corroborating the role of *PINs* in different aspects of plant development.

**Fig. 5:**
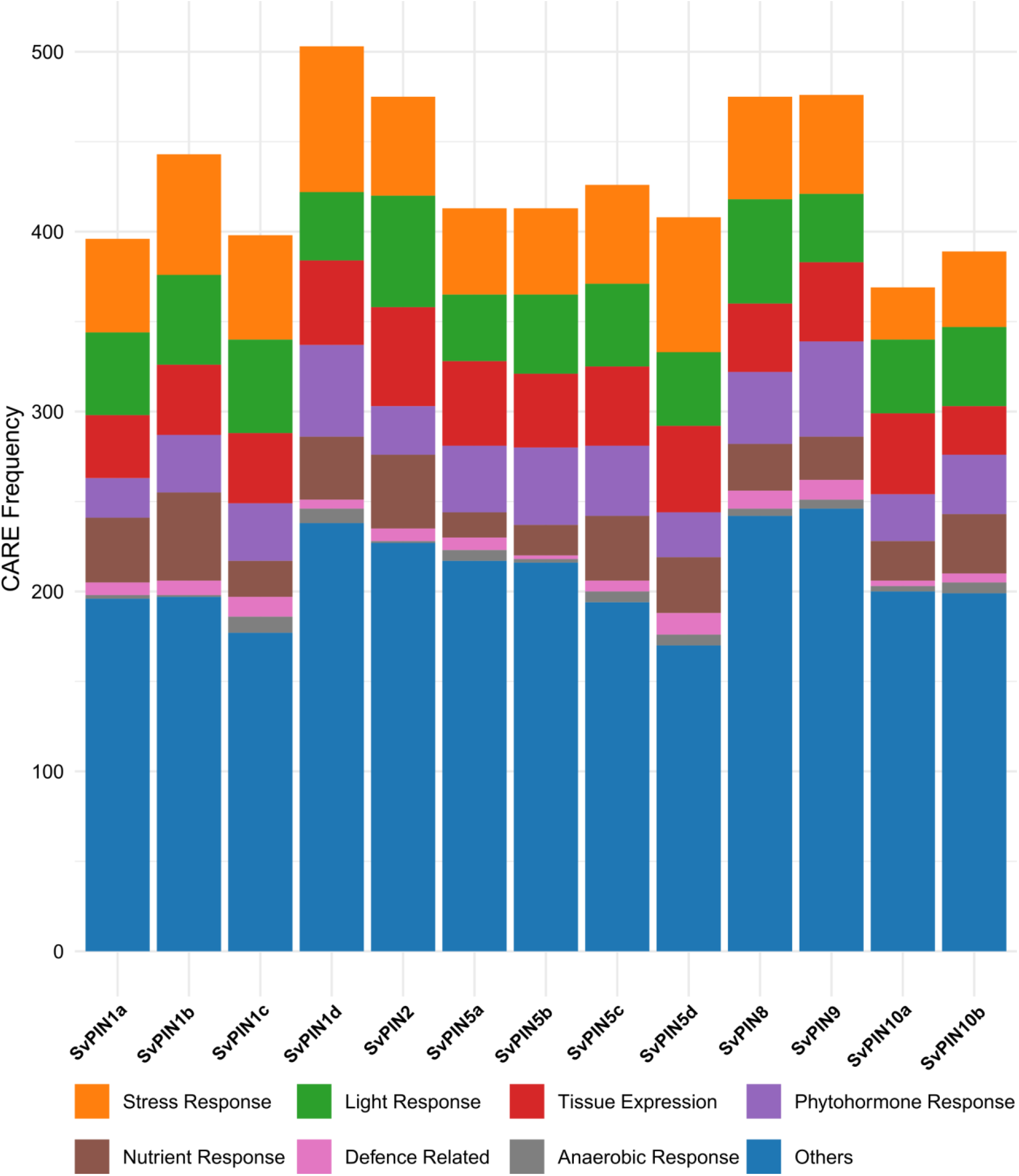
Cis acting regulatory elements (CAREs) in *SvPIN* promoters. CAREs distribution was relatively homogenous among *SvPIN* homologues. Closely related genes also displayed similarities in CARE composition in their promoters, likely due to conserved functional redundancies in the family. CAREs were grouped based on their annotated function in the NewPLACE database. A list with all identified CAREs is available in **Supplementary Table S2**.

## Discussion

Auxin is a central regulator of plant development, orchestrating processes from embryogenesis to organogenesis (Leyser 2018). One unique aspect of auxin is its polar transport, which is mediated by PIN-FORMED proteins and is crucial for vein patterning during vascular development and several other processes (Balzan et al. 2014; Wakeman and Bennett 2023). Although PINs have been extensively characterized in C_3_ model plants, such as rice and *Arabidopsis*, their role in C_4_ plants remain a significant gap in knowledge, particularly in C_4_ grasses. Investigating their role is crucial, given the importance of the specialized Kranz vascular anatomy to C_4_ photosynthetic efficiency (Sedelnikova et al. 2018). In fact, leveraging vascular development may be a key step towards introducing C_4_ metabolism into staple crops (Ermakova et al. 2020). To address this, our study provides a comprehensive characterization of the *PIN-FORMED* family throughout the development of the model C_4_ monocot *S. viridis*.

The evolutionary history of PIN proteins provides context for their functional diversity. Believed to transport auxin since the emergence of streptophyte algae, PINs were likely crucial for the colonization of terrestrial environments (Viaene et al. 2013; Bennett 2015; Bogaert et al. 2022). Within land plants, the family diversified significantly, undergoing functional innovations key to terrestrial adaptation (Zhang et al. 2020). An early divergence during the rise of the embryophytes led to the division of PINs into canonical (long, plasma membrane-localized) and non-canonical (short, ER-localized) types based on transmembrane domain structure and the length of the hydrophilic loop (Viaene et al. 2013; Viaene et al. 2014; Bennett 2015), a finding corroborated by our phylogenetic reconstruction. Further diversification was particularly important in flowering plants, where polar auxin transport and auxin maxima regulation are essential for flower development and patterning (Zhang et al. 2020). This may explain the early divergence of moss and lycophyte branches from other canonical PINs, forming a distinct “Seedless” clade in our phylogenetic tree.

We identified 13 *PIN-FORMED* genes in *S. viridis*, a number similar to other grasses like *Oryza sativa* (12 orthologues, Manna et al., 2022) and *Zea mays* (14 orthologues, Yue et al., 2015), suggesting conservation within Poaceae. This number is typically higher in monocots than in eudicots, especially grasses, likely due to duplication events (Balzan et al. 2014; O’Connor et al. 2017). For instance, the *PIN1* clade underwent triplication in grasses (Wakeman and Bennett 2023), giving rise to the *PIN1a, PIN1b* and *PIN9* clades. Furthermore, the sister clade to *PIN1*, often named *PIN11* or *SISTER-OF-PIN1* (*SoPIN1*), is a separate group absent only in Brassicaceae (O’Connor et al. 2014). While the *PIN1a/1b* duplication is widespread in grasses, our phylogenetic analysis supports the finding that the *PIN1c/1d* duplication within the *SoPIN1* clade occurs only in certain taxa, including *S. viridis, S. italica*, and *O. sativa* (O’Connor et al. 2014).

The *PIN-FORMED* family in monocots also features two exclusive clades: *PIN9* and *PIN10*. Although *PIN9* shares an origin with *PIN1*, it is a monocot-specific clade that underwent significant structural changes, resulting in a novel non-canonical group (Bennett 2015), which explains its clustering with other non-canonical *PINs* in our phylogeny. Interestingly, its presence in pineapple (*Ananas comosus*) (Zhao et al. 2021), indicates that its divergence from *PIN1* likely predates the rise of Poaceae. Similarly, the *PIN10* clade diverged significantly from its ancestral *PIN3/4/7* lineage, leading to its initial classification as a distinct group(Bennett 2015). Our phylogenetic reconstruction confirms that *PIN10* shares its most recent common ancestor with the eudicot-exclusive *PIN3/4/7* clade. *S. viridis* has two *PIN10* paralogues, *SvPIN10a* and *SvPIN10b*, and phylogenetic and collinearity analyses support a grass-specific duplication event. *PIN10* is also present in pineapple (Zhao et al. 2021) suggesting its divergence from *PIN3* predates grasses. This grass-specific diversification within *PINs* has been proposed as a driver behind developmental innovations found in Poaceae (Wakeman and Bennett 2023).

Among non-canonical *PINs*, grasses appear to have lost *PIN6* but underwent a triplication of *PIN5* (Wakeman and Bennett 2023). Some species, including *S. viridis*, possess four *PIN5* paralogues. Our collinearity analysis indicates that *PIN5a/b/c* originated from segmental duplications. The expansion of *SvPIN5* may have created functional redundancy among the paralogues. Notably, *SvPIN5a* shows a shorter transcript with fewer exons and appears to have lost two C-terminal motifs, resulting in an abnormal structure with only seven transmembrane domains. These unique characteristics, coupled with its absence of expression in all organs and calli under our experimental conditions, suggest that *SvPIN5a* may be undergoing a process of pseudogenization.

The expression patterns of *PIN* genes in *S. viridis* plants and calli provide crucial insights into their functional roles. The *PIN1* subfamily, which emerged alongside flowering plants, is known to play important roles in floral development (Zhang et al. 2020). In line with this, our observed expression patterns suggest collective roles for *SvPIN1* genes in seedling, shoot, and panicle development. PIN1a and PIN1b are known to cooperate during organ initiation, directing PAT from auxin maxima created by SoPIN1 (O’Connor et al. 2014). While these genes have distinct roles in *B. distachyon* inflorescence development, their loss-of-function mutants exhibit mild phenotypes, suggesting functional redundancy (O’Connor et al. 2014; O’Connor et al. 2017). Similar expression patterns of *SvPIN1a* and *SvPIN1b* imply conserved redundancy in *S. viridis* panicle development, though *SvPIN1a* appears to have a more prominent role in shoot and roots. Conversely, the importance of the *SoPIN1* clade is highlighted by the severe phenotypes of its mutants. *B. distachyon sopin1* knockouts resemble *Arabidopsis pin1* mutants (O’Connor et al. 2017), and rice *ospin1c-ospin1b* double mutants fail to develop inflorescences (Li et al. 2019). Additionally, *ZmPIN1d* is highly expressed in maize tassels and ears (Forestan et al. 2012). The high expression of *SvPIN1c* and *SvPIN1d* in shoots and panicles suggests similarly essential functions in *S. viridis*. Furthermore, high *SvPIN1d* expression in seedlings, coupled with CARE enrichment analysis highlighting the light-responsive element S000486, previously associated with cotyledon-specific expression in rice seedlings (Jiao et al. 2005), points to a potential role in early development.

Under high auxin, only *SvPIN1b* showed significant induction, while all *SvPIN1* genes were repressed by low auxin conditions. This aligns with findings in rice where *OsPIN1a* and *OsPIN1b* are more auxin-responsive than *OsPIN1c* and *OsPIN1d* (Li et al. 2019). While all four genes respond to auxin gradients, CARE analysis detected auxin-responsive elements exclusively in the *SvPIN1b* promoter, which could explain its stronger response. A recent study found multiple auxin-responsive CAREs in the promoters of rice *PIN-FORMED* genes (Manna et al. 2022), so their potential presence in other *SvPIN1* promoters cannot be ruled out.

The single *PIN2* orthologue in *S. viridis, SvPIN2*, displayed distinctive expression in seedlings, roots and flowering panicles. This pattern is consistent with known PIN2 functions in *Arabidopsis*. AtPIN2 mediates root gravitropism by directing PAT to root elongation zones (Balzan et al. 2014; Retzer et al. 2019) and contributes to cotyledon auxin maxima formation (Pérez-Henríquez et al. 2025). In *S. viridis, SvPIN2* expression in seedlings may relate to post-germination development, while its pattern in reproductive tissues matches *ZmPIN2* activity in maize (Forestan et al. 2012). Although PIN2 regulates rice tillering (Wang et al. 2009; Chen et al. 2012), the absence of *SvPIN2* induction in our shoot samples warrants future investigation into tillering in C_4_ grasses. Supporting its putative role in root development, the *SvPIN2* promoter contained several elements related to root and root-hair expression, such as S000314, S000482 and S000512. Furthermore, *SvPIN2* exhibited one of the strongest inductions and repressions under high and low auxin, respectively, indicating that auxin gradients are a key regulatory input for *PIN2* in *S. viridis*. This is consistent with IAA-mediated regulation of *PIN2* in *A. thaliana* (Pérez-Henríquez et al. 2025), suggesting conserved regulatory mechanisms between monocots and eudicots.

A notable divergence in expression is observed in the grass-specific *PIN10* clade. Its function has shifted from the roles of its eudicot counterpart, *PIN3*, which mediates lateral auxin transport and root tropisms (Friml et al. 2002; Zhang et al. 2013), and apical hook formation (Žádníkova et al. 2010). The apical hook absence and the differences in root architecture in monocots may explain this divergence. Instead, *SvPIN10a* and *SvPIN10b* were primarily expressed in developing and flowering panicles, with *SvPIN10b* also highly expressed in shoots during panicle initiation. This aligns with the established role of PIN10 in grass inflorescence development (Wakeman and Bennett 2023), as seen in rice and maize (Wang et al. 2009; Forestan et al. 2012). Corroborating this reproductive role, the promoter of *SvPIN10a* was found to be enriched in pollen-specific element S000254 (Hamilton et al. 1998).

When it comes to *PIN5*, the absence of *SvPIN5a* expression, combined with its truncated transcript and anomalous protein structure missing transmembrane domains, strongly supports the pseudogenization hypothesis, likely stemming from redundancy within the subfamily. Usually, PIN-FORMED proteins display five transmembrane helices on both sides of the hydrophilic loop (Ung et al. 2022). The missing three helices on the C-terminus of SvPIN5a correlate with the missing motifs in our motif analysis and may indicate that this is a non-functional protein. The remaining three *SvPIN5* paralogues were expressed mainly in leaves and reproductive tissues, contrasting with the role of *AtPIN5* in root development (Mravec et al. 2009; Ganguly et al. 2014). The high expression of *SvPIN5b* in leaves suggests a potential involvement in monocot vein patterning possibly by coordinating intracellular auxin transport in association with PIN1 (Sawchuk et al. 2013). This parallels the expression of *OsPIN5a* in rice (Wang et al. 2009) and result warrants further investigation into its role in C4 grass vascular development. Similarly, the expression of *SvPIN5c* and *SvPIN5d* in reproductive tissues mirrors *ZmPIN5c* expression in developing maize kernels (Forestan et al. 2012), suggesting an involvement in seed development. This is supported by the enrichment of embryogenesis-associated element S000042 (Chandrasekharan et al. 2003) in the *SvPIN5d* promoter, which may explain its peak expression during late reproductive stages when the first seeds start to develop.

Finally, the expression of the remaining *PINs* also reveals functional insights. *SvPIN8* was highly expressed in shoots and inflorescences, matching expression patterns in rice and maize (Matthes et al. 2019), but contrasting with *Arabidopsis*, where *AtPIN8* is predominantly expressed in pollen grains and during the formation of the pollen tube, while showing very low expression in the inflorescence itself (Ding et al. 2012; Bosco et al. 2012). This indicates a departure from the role of PIN8 in eudicots, since expression in grasses seems to be greatly reduced in pollen grains (Wakeman and Bennett 2023). *SvPIN9* showed elevated expression in mature roots, a pattern consistent with observations in rice and maize plants (Wang et al. 2009; Forestan and Varotto 2012). The increase in *SvPIN9* expression between stages 1.70 and 5.50 coincides with the development of a larger root system, suggesting a potential role in lateral root formation. Additionally, given that PIN9 has been implicated in rice tillering, especially in the waterlogged conditions of flooded paddy fields (Nguyen et al. 2018; Hou et al. 2021), this further highlights the importance of studying tillering in C_4_ grasses. Collectively, these expression profiles not only affirm the conserved roles of several *PIN* orthologues in fundamental developmental processes but also highlight clade-specific functional innovations within grasses, providing a foundational framework for understanding auxin transport in the C_4_ context.

## Conclusion

Our analysis of the *PIN-FORMED* family in *Setaria viridis* reveals key evolutionary and functional insights into auxin transport in C_4_ grasses. We identified 13 *SvPIN* genes and uncovered lineage-specific duplications, underscoring a history of functional diversification. Distinct expression patterns indicate specialized, with *SvPIN1* and *SvPIN10* in panicle development, while *SvPIN2* and *SvPIN9* underly root development. Notable findings like the pseudogenization of *SvPIN5a* and the distinct hormonal responsiveness of *SvPIN1b* and *SvPIN2* highlight a complex regulatory landscape. Our findings lay a foundational groundwork for future investigations aiming to leverage auxin transport pathways to improve C_4_ crop architecture and contribute to more sustainable agriculture practices.

## Author Contribution Statement

All authors contributed to the study conception and design. Material preparation, data collection and analysis were performed by JMFE, JTL and JDFG. The first draft of the manuscript was written by JMFE, and all subsequent versions of the manuscript were reviewed by all authors. Funding was secured by MAF. All authors read and approved of the final manuscript.

## Data Availability

All data generated and analyzed during this study is included in this published article and its supplementary files.

## Statements and Declarations

