## Supplementary Material for "The Evolution and Developmental Expression Profile of the *PIN-FORMED* Family in *Setaria viridis*"

*Corresponding author

ORCID and e-mail

**Supplementary Material**

**Supplementary Table S1:** CARE list.

**Supplementary Table S2:** CARE enrichment analysis.

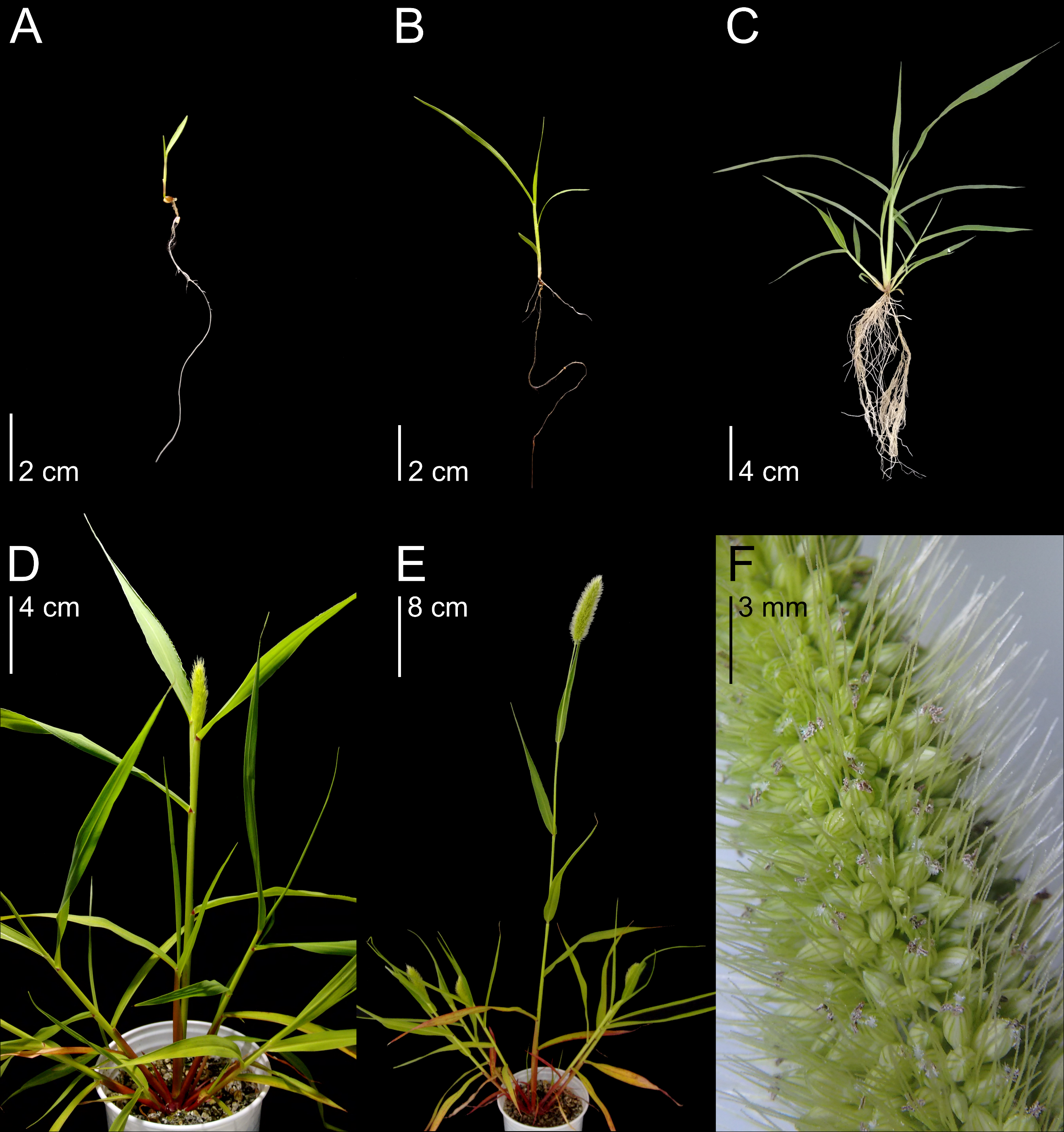

**Supplementary Fig. S1: *Setaria viridis* developmental stages analyzed through qPCR. (A)** Stage 1.10, 8 days after imbibition (DAI). Seedling after full expansion of the first leaf. **(B)** Stage 1.30, 15 DAI. Full expansion of the third leaf. Initiation of lateral root growth. **(C)** Stage 1.70, 24 DAI. Full expansion of the seventh leaf. Development of tillers and significant root architecture expansion. **(D)** Stage 5.50, 33 DAI. Partial panicle emergence, marking the start of the reproductive phase. **(E)** Stage 7.10, 42 DAI. Fully developed panicle and first visible diaspores. **(F)** Detail of the 7.10 panicle, with developed diaspores.

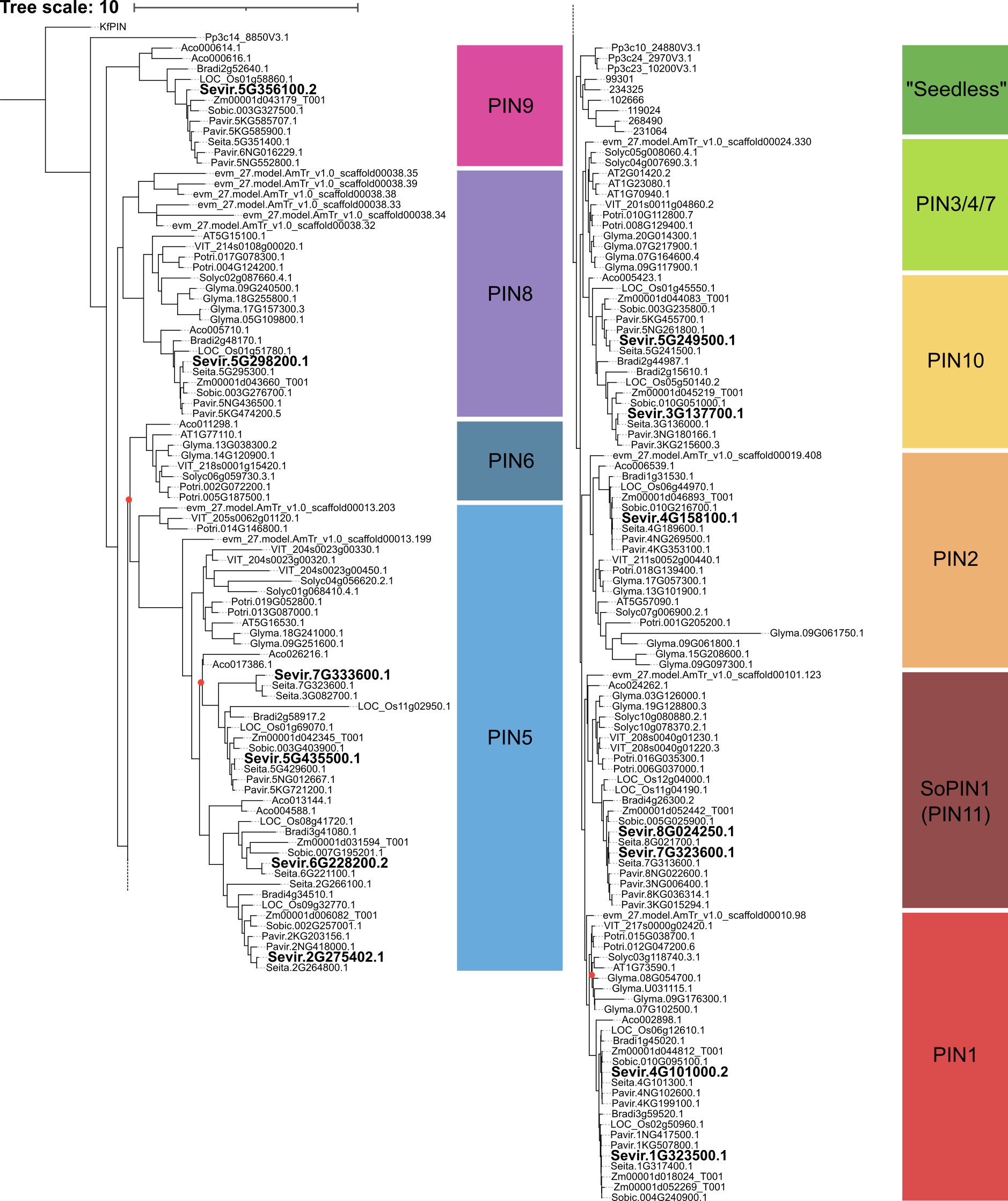

**Supplementary Fig. S2:** **Maximum-likelihood phylogenetic tree with branch lengths.** *Setaria viridis* orthologues are highlighted in bold. Branches with support lower than 70% are marked with a red dot. The charophyte *K. flaccidum PIN* orthologue (*KfPIN*) was used as the outgroup.

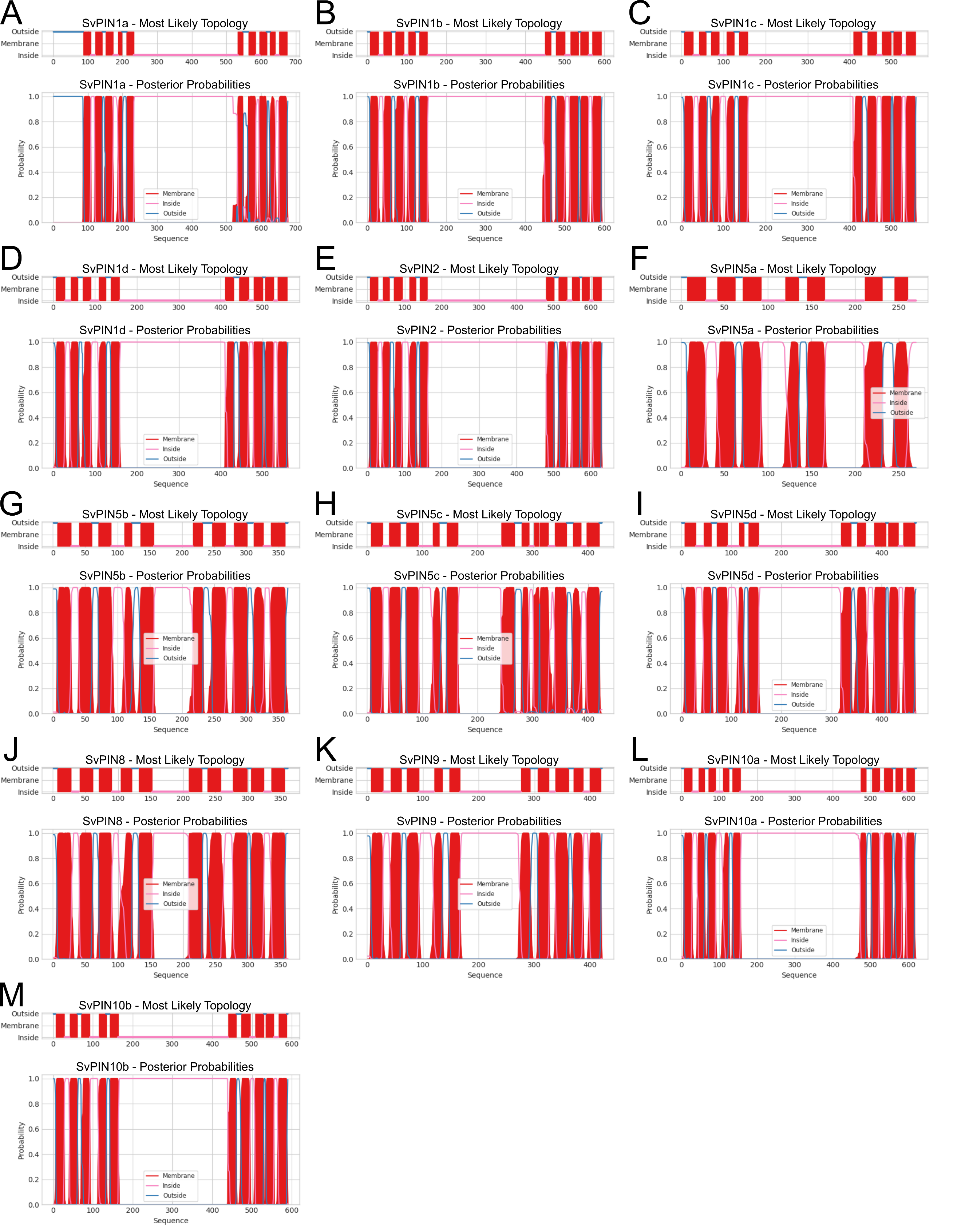

**Supplementary Fig. S3: Transmembrane domain prediction of *Setaria viridis* PIN homologues.** Canonical and non-canonical proteins display differently sized central hydrophilic loops, whereas canonical PINs have longer loops than non-canonical PINs.

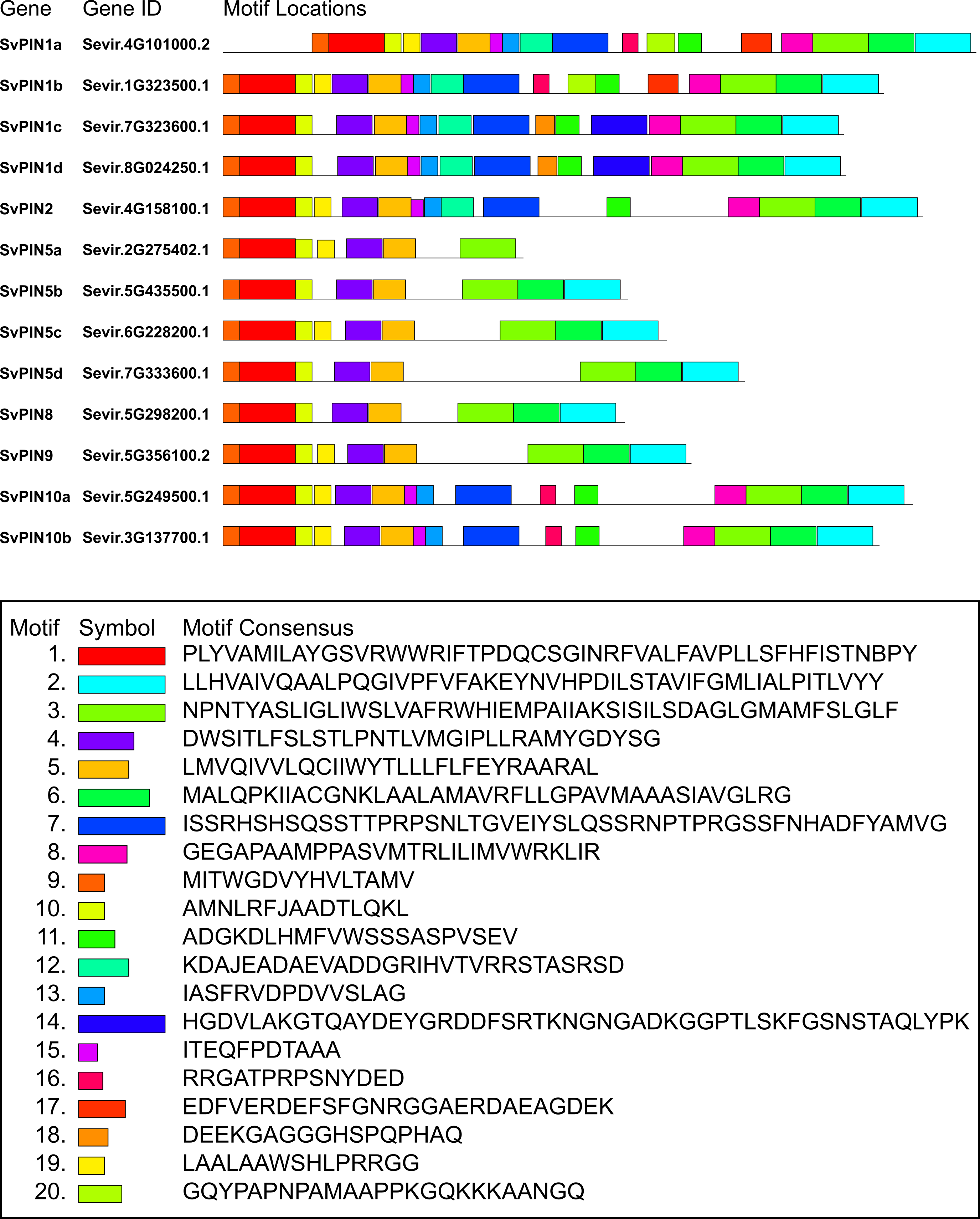

**Supplementary Fig. S4: Conserved protein motifs and consensus sequences.** All PINs showcase highly conserved motif compositions, with the same N- and C-terminal motifs being present in almost all 13 proteins. SvPIN5a, however, lacks the last two conserved motifs (numbers 3 and 6), due to its shorter transcript, resulting in an abnormal protein with only seven transmembrane domains.
